# VOLARE: Visual analysis of disease-associated microbiome-immune system interplay

**DOI:** 10.1101/431379

**Authors:** Janet C. Siebert, Charles Preston Neff, Jennifer M. Schneider, EmiLie H. Regner, Neha Ohri, Kristine A. Kuhn, Brent E. Palmer, Catherine A. Lozupone, Carsten Görg

**Affiliations:** Computational Bioscience Program, University of Colorado Anschutz Medical Campus, Aurora, CO 80045, USA; CytoAnalytics, Denver, CO 80113; Department of Medicine, University of Colorado Anschutz Medical Campus, Aurora, CO 80045, USA

## Abstract

**Background:** Relationships between specific microbes and proper immune system development, composition, and function have been reported in a number of studies. However, researchers have discovered only a fraction of the likely relationships. High-dimensional “omic” methodologies such as 16S ribosomal RNA (rRNA) sequencing and Time-of-flight mass cytometry (CyTOF) immunophenotyping generate data that support generation of hypotheses, with the potential to identify additional relationships at a level of granularity ripe for further experimentation. Pairwise linear regressions between microbial and host immune features is one approach for quantifying relationships between “omes”, and the differences in these relationships across study cohorts or arms. This approach yields a top table of candidate results. However, the top table alone lacks the detail that domain experts need to vet candidate results for follow-up experiments.

**Results:** To support this vetting, we developed VOLARE (Visualization Of LineAr Regression Elements), a web application that integrates a searchable top table, small in-line graphs illustrating the fitted models, a network summarizing the top table, and on-demand detailed regression plots showing full sample-level detail. We applied VOLARE to three case studies—microbiome:cytokine data from fecal samples in HIV, microbiome:cytokine data in inflammatory bowel disease and spondyloarthritis, and microbiome:immune cell data from gut biopsies in HIV. We present both patient-specific phenomena and relationships that differ by disease state. We also analyzed interaction data from system logs to characterize usage scenarios. This log analysis revealed that, in using VOLARE, domain experts frequently generated detailed regression plots, suggesting that this detail aids the vetting of results.

**Conclusions:** Systematically integrating microbe:immune cell readouts through pairwise linear regressions and presenting the top table in an interactive environment supports the vetting of results for scientific relevance. VOLARE allows domain experts to control the analysis of their results, screening dozens of candidate relationships with ease. This interactive environment transcends the limitations of a static top table.

## 1 Background

“Omic” approaches such as transcriptomics, metabolomics, and mass cytometry allow researchers to measure hundreds to thousands of analytes. However, data from a single ome may lack rich functional insight (1) or may miss signals that are present in another ome (2). Thus, multi-omic studies are increasingly common (3,4), offering the potential to formulate progressively more comprehensive perspectives on biological processes (5–8). Multi-omic studies may interrogate closely related omes, such as genes and their methylation (3), or more disparate omes, such as the gut microbiome and immune cell subsets (9). Among the challenges of such studies are analyzing the data to identify specific cross-omic patterns. As an example of one such pattern, *Bacteroides fragilis* induces regulatory T cells to produce IL-10, conferring protection from inflammation in mouse models (10). Aberrations in both the gut microbiome and the immune system have been associated with diseases including inflammatory bowel disease (11), type 1 diabetes (12), asthma (13), multiple sclerosis (14), rheumatoid arthritis (15), and HIV (16,17); and in responses to immunotherapy (9,18,19). However, researchers have discovered only a fraction of the underlying relationships and their associations with disease. Identification of cross-omic patterns in multi-omic data offers the potential to identify additional candidate relationships at a level of granularity ripe for further experimentation. Furthermore, these relationships can connect an analyte of interest from one ome to unfamiliar analytes in another ome. For example, an immunologist studying patient responses to an immunotherapy that blocks an inhibitory receptor, such as programmed cell death 1 (PD-1), might be interested in commensal microbes that are associated with cell populations that express PD-1 (20). The ability to identify cross-omic relationships is of interest both to a single researcher incorporating new omic technologies into his or her studies, and to a cross-disciplinary research team.

One approach to identifying cross-omic relationships is to systematically compare all of the analytes in one ome to all of the analytes in another ome, using either correlation (9) or regression techniques (18,21). The resulting data can be presented as a heat map of correlation coefficients (9,22) or p-values (18). Alternatively, we can focus on a “top table” of statistically significant associations, similar to those generated in the analysis of gene expression data (23). This top table lists which microbes are associated with which immune cells (Supplemental Table 1). However, the top table alone lacks the detail that a researcher needs to prioritize results for follow-up laboratory experiments. In the case of cross-omic regressions that account for difference in disease state, each row in a top table represents a complex relationship for each pair of analytes, not well captured by a test statistic alone. To support visual analysis of these relationships, and help researchers prioritize results for follow-up, we developed a novel web application called VOLARE, (Visualization Of LineAr Regression Elements). VOLARE provides a visual encoding of the top table and associated regression elements, leveraging existing visualization techniques. We extend the top table, a fundamental tool of single-omic analysis, to two omes. We enrich it with small in-line graphs of the fitted regression models, from which the researcher can drill down to detailed regression plots illustrating both the fitted model and sample-level detail. The table itself (or a subset thereof) is summarized by an interactive network, with analytes represented as nodes and relationships as edges. This interactive environment supports visual data analysis and transcends the limitations of a static top table. This approach is broadly applicable to studies that include data from two or more high-throughput assays, such as microbe:metabolome, microbiome:proteome, and RNA-Seq:immune repertoire.

The overall goal of our approach is to support vetting of results for scientific relevance. Through structured interviews and ongoing collaboration with domain experts, we identified the following tasks associated with the vetting process: (1) explore relationships between an analyte of interest and associated analytes in the other ome, thereby borrowing information from one domain to better understand another; (2) discover relationships that differ across disease state (e.g. HIV+ and HIV-); (3) assess credibility of the fitted model, including goodness of fit, the presence or absence of outliers, and the magnitude and dynamic range of the readouts for each analyte; (4) compare detailed regression plots across several pairs of analytes; and (5) identify highly connected “hub” analytes, such as a particular microbe related to a number of immune cell subsets. To illustrate generalizability, we applied VOLARE to three case studies: microbe:cytokine data from fecal samples; microbe:cytokine data, with cytokines produced by ex-vivo mitogen stimulation of intraepithelial lymphocytes; and microbe:immune cell data from gut mucosal biopsies.

## 2 Methods

### Architecture and workflow

Figure 1 illustrates the VOLARE architecture and workflow, which consists of preparing data for the web application, and using the web application itself. We used the R statistical programming environment to prepare data for VOLARE, and implemented the web application in HTML, JavaScript, and the D3 library (24). The data preparation processes generate regression results from merged assay data, and format the top table results and underlying detail into a JavaScript Object Notation (JSON) file with the jsonlite library (25). The domain expert then loads this JSON file into VOLARE for visual analysis. Our architecture recognizes that the processes of merging multi-omic data for analysis, performing thousands of pairwise linear regressions and generating an associated top table, and analyzing the results in the top table are distinct and often performed by people with different areas of expertise. Because it reflects a fundamental separation of concerns, this architecture can be applied to a variety of omes. It also results in a visual analysis environment that is responsive to user input, since the computationally intensive calculations are performed upstream of visualization. We provide representative example scripts for performing regressions and formatting the data at https://sourceforge.net/projects/cytomelodics. Source code, a tutorial, and example input files are also available at https://sourceforge.net/projects/cytomelodics. A hosted version of VOLARE is available at http://aasix.cytoanalytics.com/volare/, and includes a link to the JSON file used for Case Study 3, discussed below.

**Figure 1.**
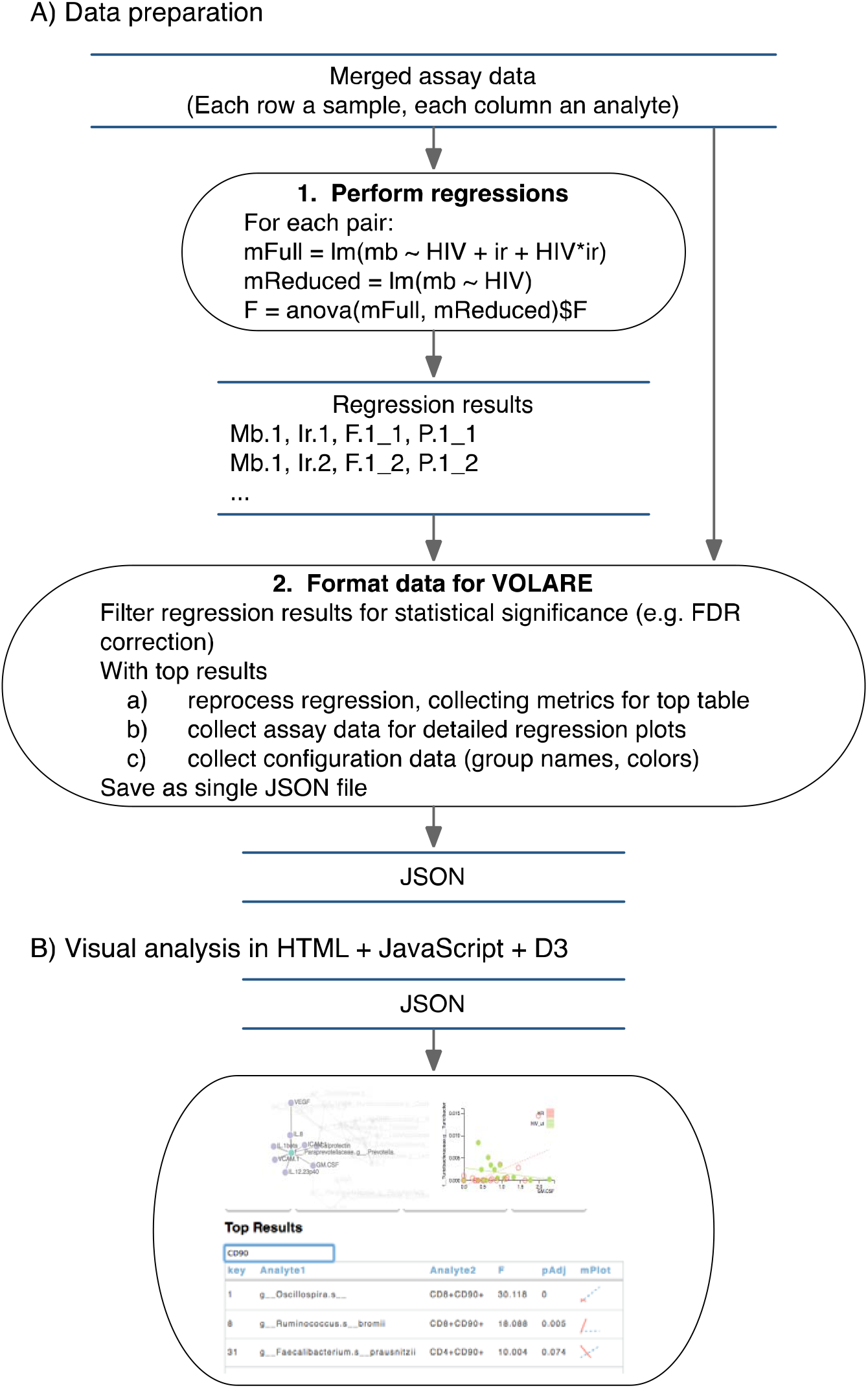
VOLARE architecture and workflow. The architecture reflects a separation of concerns between data preparation and visual analysis. Blue horizontal parallel lines represent data files. Black ovals represent processes. A) Data preparation is performed in R. Given a file of merged assay data, all pairwise regressions are calculated and recorded, with Mb.1 and Ir.1 representing the first microbe and first immune readout respectively. F.1_1 and P.1_1 represent the F statistic and p-value from the linear model using Mb.1 and Ir.1. Second, data is formatted for VOLARE. Regression results are filtered for statistical significance. These top table results are reprocessed to collect additional details needed for visualization (top table of relationships and associated metrics, underlying data, and configuration data for the web application), which are saved in a JSON file. B) Visual analysis is performed with a JavaScript web application.

### Regression models

To address the question, “is the relationship between any particular microbial taxa (Mb) and any particular immune readout (IR) different based on cohort?” we used a partial F-test comparing the linear regression model, Mb ~ Cohort + IR + Cohort x IR to a reduced model, Mb ~ Cohort. This tests whether the full model has more explanatory value than does the reduced model. Specific cohorts and immune readouts are discussed in the context of the case studies.

### Visual design

Figure 2 illustrates the VOLARE visual analysis interface. Since the top table is a fundamental element of omics analysis, we built VOLARE around the table. To support **Task 1** (explore relationships between an analyte of interest and associated analytes in the other ome), we added an interactive filter function to the top table. When the domain expert enters a microbe or immune marker, the table automatically displays only those relationships that match the search phrase. While we could have represented the top table as a matrix or heat map, the textual and numeric details of the table are essential to communicate the results of the statistical analysis. Furthermore, the VOLARE top table displays all of the columns that were included in the top table structure in the JSON file. These columns can include mean values or observed ranges of each analyte, p-values of cohort-immune response interaction terms (top table in Figure 5C), or influence measures (top tables in Figures 6A and 6B), thereby placing additional derived data in context.

**Figure 2.**
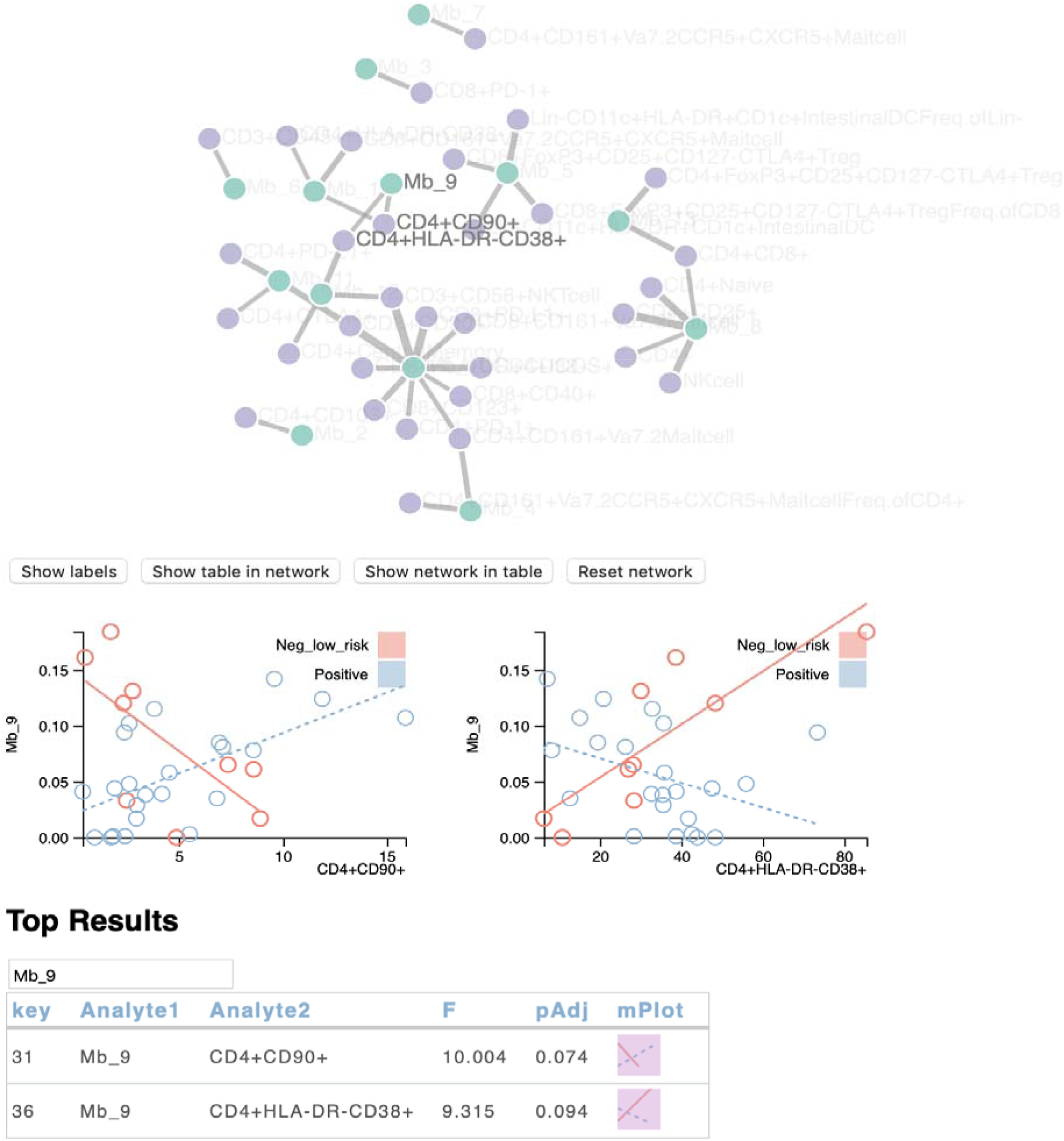
VOLARE screenshot: Network at the top, two detailed regression plots below, and top table at the bottom. Buttons add labels to the nodes, synchronize the table with the network, or synchronize the network with the table. The top table can be filtered by typing text to match. The table contains one row for each relationship, listing the analytes that comprise the relationship, the test statistic (in this case F), an adjusted p-value (pAdj), and a small plot illustrating the fitted model (mPlot). Clicking on an mPlot generates the corresponding detailed graph. In each detailed plot, the x-axis represents the immune cell population (measured in percent of parent population) while the y-axis represents the microbial taxa (measured in relative abundance, in the range from 0 to 1). Each point represents the values for one sample from one person. Points are color coded to represent the cohort to which the corresponding person belongs. Lines represent the fitted regression model for each cohort. The closer the points are to the line, the better the model.

To support **Task 2** (discover relationships that differ across disease state), we added small graphs of the fitted regression model, inspired by Tufte’s sparklines (26). We call this embedded graphic the microplot or “mPlot.” The graphic encoding of this derived data enables the domain expert to scan the table and quickly assess what analytes are involved in what sorts of relationships.

As such, it also functions as a small multiple display. The mPlot illustrates the regression model using line tilt, line length, and color. While the same data could be represented by numeric values for slope, such an encoding would be less conducive to visual analysis. Furthermore, the magnitude of the analyte readouts (and thus the slopes) can vary widely across the data set. The mPlot normalizes the magnitudes by plotting the relationship in a consistently sized glyph, regardless of the magnitude. Figure 3 provides three different mPlot examples, with different interpretations. Figure 3A illustrates a relationship in which the microbe and immune readout are associated in one cohort but not the other, possibly because the microbe is not present in one of the cohorts. Figure 3B illustrates a positive association in one cohort and negative association in the other, which might suggest differing biological mechanisms in health and disease. Figure 3C illustrates a much smaller dynamic range of both analytes in one cohort than the other. While this could be driven by a single outlier, it also could indicate truly different ranges in both analytes across the two cohorts. Thus, even though the mPlot provides a valuable glimpse of the relationship between the analytes, underlying detail is required to fully vet the relationship.

**Figure 3.**
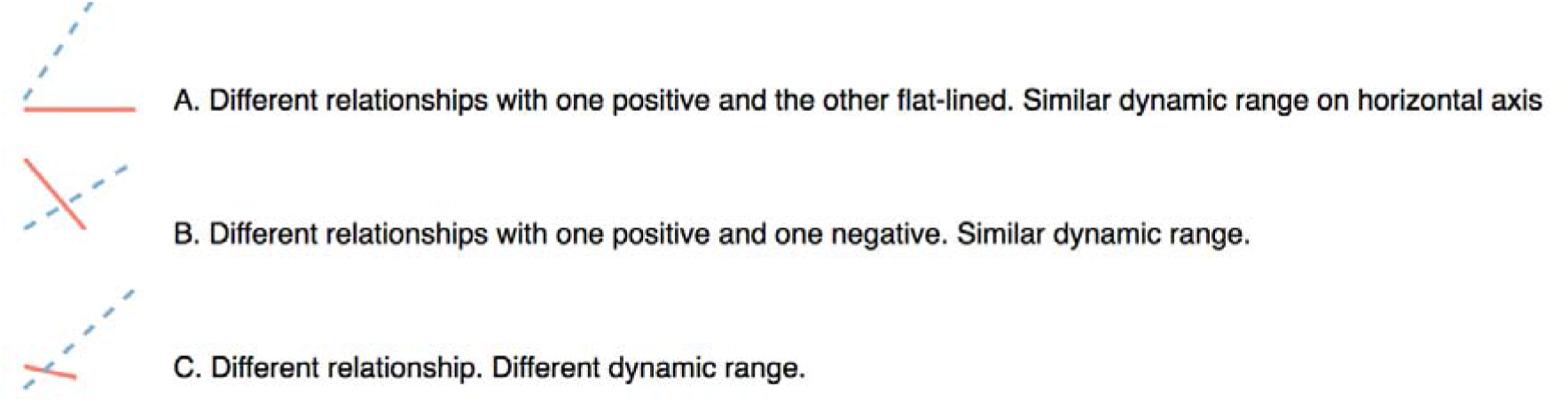
mPlot examples. Solid and dotted lines represent different cohorts. The vertical axis represents microbe relative abundance, while the horizontal axis represents the immune readout. Three examples illustrate the different relationships that can be encapsulated in the sparkline-inspired mPlot. (A). A relationship between the microbe and the immune readout exists in one cohort but not the other which might suggest that the microbe is absent in the “flat line” group. (B). Differences in relationship between the microbe and immune readout across the two cohorts might suggest biological differences across the cohorts. (C). The difference in dynamic range across the cohorts might suggest that the relationship captured by the longer line is driven by an outlier, with high values in both analytes.

To support **Task 3** (assess credibility of the fitted model, including goodness of fit, the presence or absence of outliers, and the magnitude and dynamic range of the readouts for each analyte) and **Task 4** (compare detailed regression plots across several pairs of analytes), we provide a detailed regression plot in response to clicking the mPlot. Multiple plots can be juxtaposed in the same view to support comparison. This detailed plot illustrates each data point, colored to indicate disease status, and the corresponding regression fit. The encoding of a detailed regression plot necessary to convey statistical detail aligns well with best practices of visual encoding. Each point is grounded in a common two-dimensional space, color indicates groups, and tilt captures the fitted model (27).

To support **Task 5** (identify highly connected “hub” analytes), we present a network that summarizes the relationships in the top table. Each node represents an analyte with color encoding the assay (e.g., purple = immune cell subset, green = microbe), while each edge indicates a relationship between two analytes, i.e. a row in the top table. This is an efficient use of screen real estate in which each analyte from the top table appears only once, with relationships captured by edges. Alternatively, we could have summarized the table with a histogram of analyte degree, but this would not have included the relationships between analytes. Taken together, these encodings support the Shneiderman mantra of overview first, zoom and filter, then details on demand (28). The network and top table provide the overview. The mPlots provide a pre-zoomed representation. The top table itself can be filtered, and the detailed plots are available on demand.

### Biological methods and materials

Biological methods and materials are presented in Supplemental File 1.

## 3 Results

We applied VOLARE to data sets from three different studies. The first case study interrogates microbiome:cytokine relationships in fecal samples from HIV-negative high risk individuals and HIV-negative low risk individuals. The second case study uses published data, and identifies new findings in fecal microbiome:cytokine relationships in patients with spondyloarthritis, Crohn’s disease, ulcerative colitis, and healthy controls (29). The third case study considers microbiome:immune cell relationships in gut biopsies in HIV-positive and HIV-negative participants. In all three cases, we examine relationships between microbial taxa and immune readouts. Since one of our goals is to identify a “reasonable” number of candidate results for vetting (about 30 to 100), we use a different cutoff for inclusion in the top table in each case. We generate top tables with different sets of metrics according to study design and research questions. In one case, we apply a square root transformation on cytokine data. In another case, we illustrate influential observations by encoding an influence metric in the size of the circles in the detailed plot. Finally, we characterize user interaction with VOLARE by analyzing server logs.

### Case study 1: Microbiome:cytokine relationships in HIV

Fecal samples provide a non-invasive source of microbiota and proteins generated by immune cells. Here, we describe an unpublished study using such samples, and analysis of the resulting data using VOLARE. Fecal samples were collected from study participants who were HIV negative high risk (HR; men having sex with men, n=17) or low risk (LR, n=18). High risk individuals engage in behaviors that put them at increased risk for acquisition of HIV. Fecal samples were analyzed by 16S rRNA sequencing to identify microbes and by ELISA to identify a combination of cytokines and growth factors; hereafter, called cytokines. To compare microbes to cytokines, we combined data for 35 study participants into a single file consisting of 43 microbial genera with non-zero relative abundance values for at least 17 of 35 samples and 17 cytokines. We fitted 731 (43 × 17) linear regression models of the form Mb ~ Cohort + Cytokine + Cohort x Cytokine and compared those results to those from a reduced model, Mb ~ Cohort using a partial F-test. We surfaced the 58 pairs with an unadjusted p < 0.05 for exploration in VOLARE.

Figure 4 illustrates analysis tasks as defined in the *Background* section in the context of this case study. At a high level, the domain expert identifies an analyte of interest based on prior knowledge, network community, or mPlot trends, filtering the table to display the rows that include this analyte. Inspecting the detailed plots may in turn lead to the identification of a new analyte of interest. First, we searched for a specific microbe of interest, “Mb_6.” The filtered table has only one row, showing that Mb_6 is associated with IL-1α (Figure 4A, **Task 1**). In this case, there is a strong negative association between the bacteria and IL-1α for the low risk group (LR in blue), while the relationship between Mb_6 and IL-1α for the high-risk group is relatively flat (HR in red; **Task 2**). Clicking on the mPlot generates the detailed plot. Here, we observed that several people in the high-risk group have high levels of IL-1α, represented by the rightmost points with values around 1,600 and 1,800 pg/ml (Figure 4B, **Task 3**). Thus, IL-1α is of interest. To see if other bacteria are associated with these high IL-1α values, we searched the top table for IL-1α (Figure 4C, Task 1), drilling down on the several detailed plots (Figure 4D, **Task 4**). We visualized the relationships between microbial taxa and IL-1α in a network graph (Figure 4E, Task 5). Overall, we observed that the IL-1α outliers were associated with high levels of Mb_8, but not with high levels of Mb_12. We speculated that Mb_8 was driving an IL-1α immune response, and considered an *in vitro* experiment to recapitulate this association in cells from other study participants.

**Figure 4.**
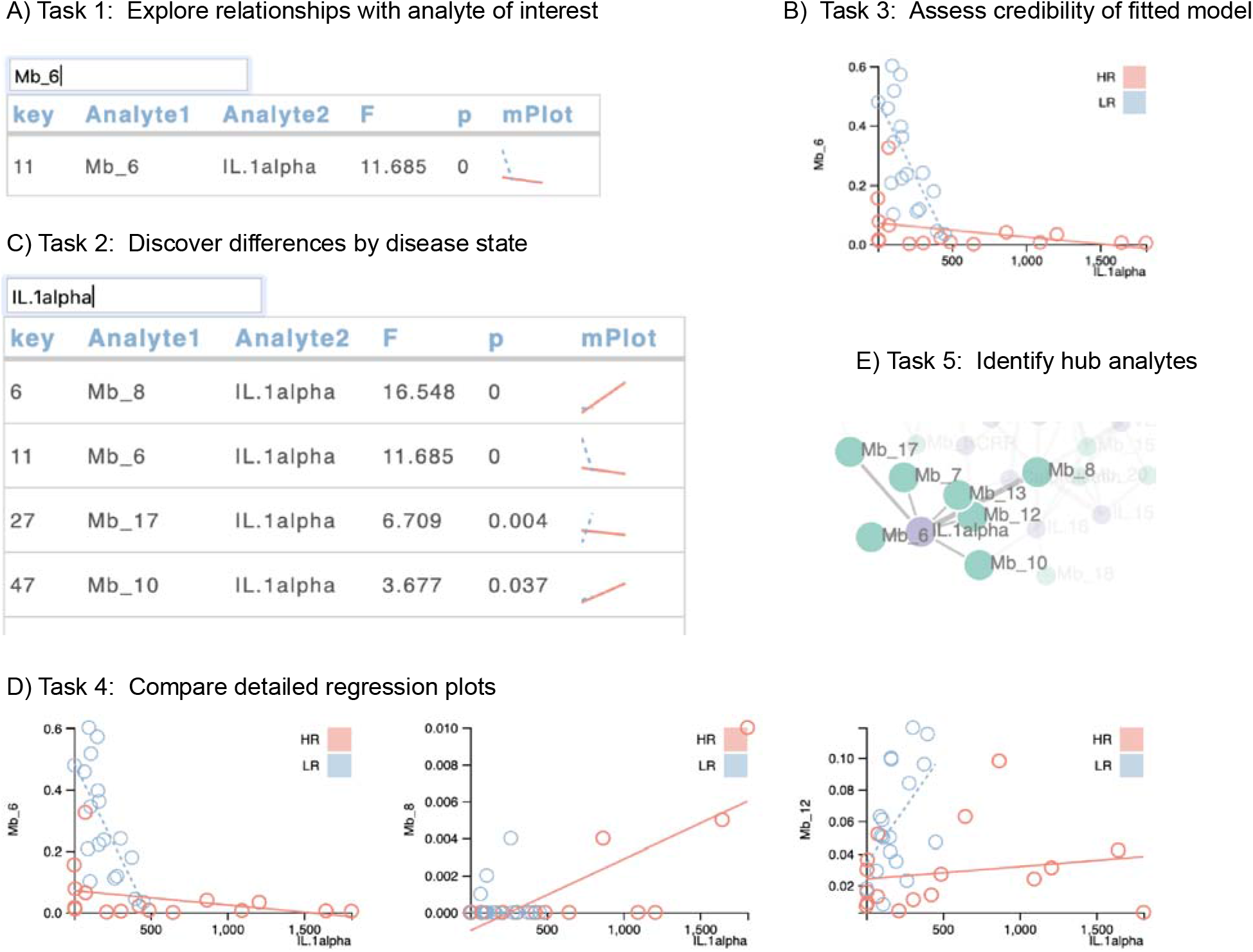
Case study 1 mapped to tasks. (A) To explore relationships, we searched for a microbe of interest in the top table, and then (B) generated a detailed regression plot to assess credibility of the fitted model. The x-axis represents the cytokine data (measured in pg/ml) while the y-axis represents the microbial taxa (measured in relative abundance, in the range from 0 to 1). Each point represents the values for one sample from one person. Points are color coded to represent the cohort to which the corresponding person belongs. Lines represent the fitted regression model for each cohort. The closer the points are to the line, the better the model. The relatively large dynamic range of the values for IL.1alpha make it an analyte of interest. (C) Partial results of the search for IL.1alpha. mPlots allow us to discover differences by disease state. (D) Comparing detailed regression plots, we obsered that high values of IL.1alpha are associated with relatively high levels of Mb_8 but not Mb_12. (E) The ability to show our IL. 1alpha table in the network illustrates that IL. 1alpha is a a hub connected to 7 proteins.

### Case study 2: Microbiome:immune cell relationships in inflammatory bowel disease and spondyloarthritis

Previously, Regner et al. reported on relationships between the gut microbiome and cytokines produced by mitogen-stimulated intraepithelial lymphocytes (IEL) in patients with spondyloarthritis (SpA), Crohn’s disease (CD), ulcerative colitis (UC), and healthy controls (HC) (30). Among other results, Regner identified elevated levels of TNFα in patients with SpA and CD. To compare gut microbiome to cytokines produced *ex vivo* by mitogen-stimulated IEL, we combined data for 37 study participants (across 4 cohorts) into a single file consisting of 70 microbial taxa and 6 cytokines. Cytokine values were square root transformed to compress the dynamic range of the data and dampen the effect of very high readings. We fitted 420 (70 × 6) linear regression models of the form Mb ~ Cohort + Cytokine + Cohort x Cytokine and compared those results to a reduced model, Mb ~ Cohort, using a partial F-test, surfacing 32 pairs with an FDR adjusted p-value < 0.05 for exploration. We included the p-values for the cohort:cytokine interaction terms in the VOLARE top table.

To follow up on the TNFα finding, we focused on relationships between microbes and TNFα **Task 1**), observing a strong relationship with Bacteroidales/S24-7 in both CD and SpA (Figure 5A, **Task 3**). Next, we searched for other relationships with Bacteroidales/S24-7, finding a relationship with IL-6 (Figure 5B, **Task 3**). While the detailed plot suggests that this relationship was driven by a single outlier, high for both IL-6 and S24-7, we wondered if this patient had relatively high levels for other microbes. Thus, we searched for IL-6 (Figure 5C, **Task 1**), finding four other microbes (Rikenellaceae/RC9-gut-group, Porphyromonadaceae/Odoribacter, Bacteria/Candidate-division-TM7, and Clostridiales/Ruminococcaceae) in which the microbe:IL-6 relationship for this patient was also aberrant, as shown in the detailed plots in Figure 5D (**Task 4**). These results suggest that there might be patient-level patterns of microbe:cytokine relationships associated with disease state.

**Figure 5.**
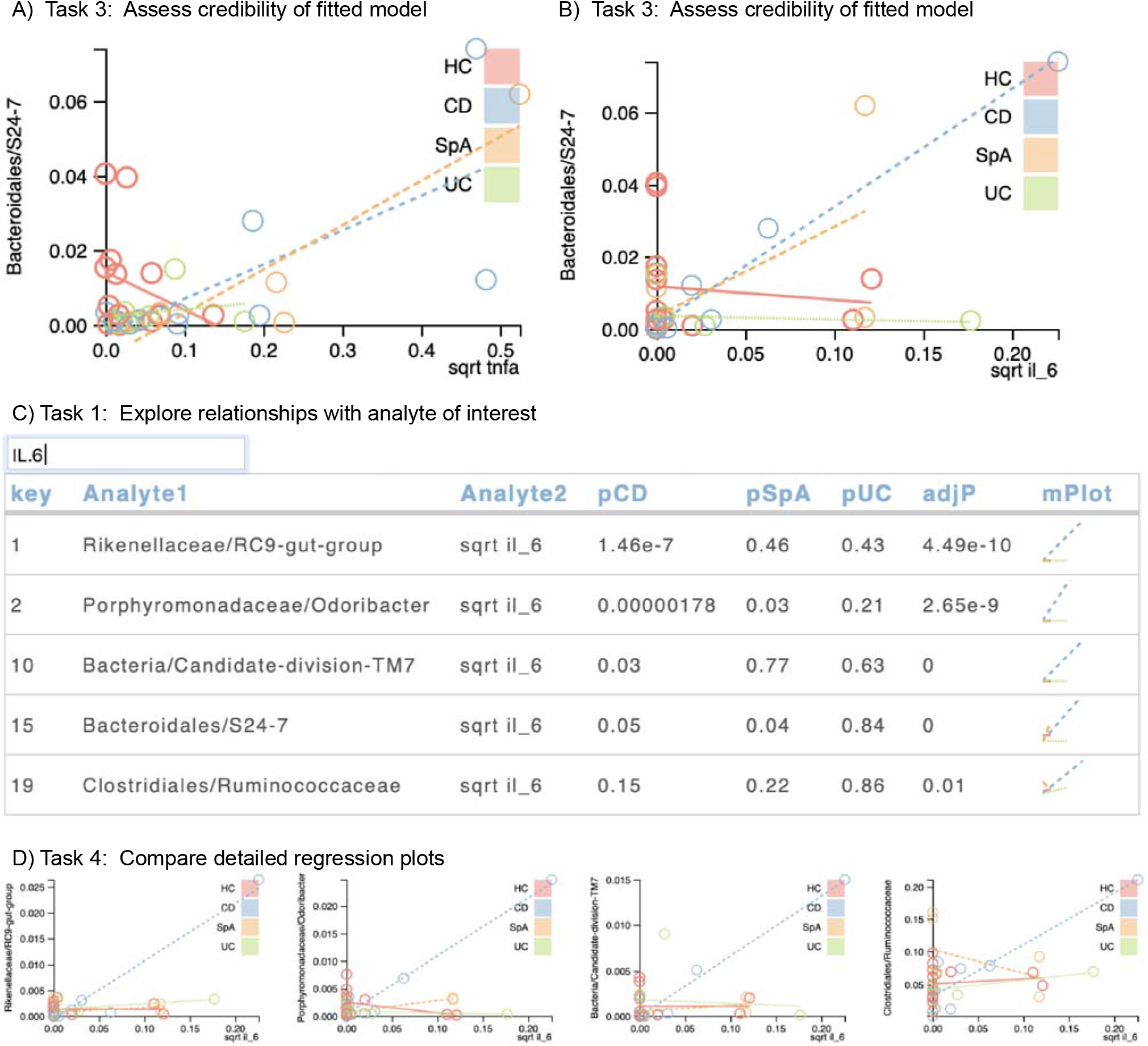
A). TNFa and Bacteroidales/S24-7 are positively associated in both SpA and CD. B) IL-6 and Bacteroidales S24-7 are also positively associated in SpA and CD, with one CD patient showing high levels of both analytes. C) A subsequent exploration of IL-6 shows positive associations between IL-6 and four other microbial taxa. In this example, the top table includes p-values for the interaction terms for each of three cohorts (CD, SpA, and UC) with respect to the reference group of healthy controls (HC). D) The detailed plot shows that the patient with the highest IL-6 values is also relatively high in four other microbial taxa.

### Case study 3: Microbiome:immune cell relationships in HIV

We considered the interplay between the microbiome and immune cell repertoire in gut biopsies of 18 volunteers, half of whom were HIV+ and half HIV-. We combined data into a single file consisting of 54 microbial genera with non-zero relative abundance values for at least 9 samples and 103 immune cell subsets. We fitted 5,562 (54 × 103) linear regression models of the form Mb ~ Cohort + Immune cell + Cohort x Immune cell and compared those results to a reduced model, Mb ~ Cohort, using a partial F-test. We surfaced 78 results with an FDR adjusted p-value < 0.1. Through visual analysis, we identified several cases in which a microbe was associated with an immune cell subset in health (HIV-) but not in disease (HIV+). As an example, *Bactemides* genus positively associated with CD4+FOXP3+ and CD4+HLA-DR+CD38- T cell populations (Figure 6A) in samples from HIV- participants. This FOXP3+ association is concordant with prior work that shows an increase in regulatory T cells in response to stimulation with *Bacteroides fragilis* lysates (10). Prior work also shows an induction of CD4+HLA-DR+CD38+ T cells in response to stimulation with whole fecal bacterial communities (30). While we have not previously focused on HLA-DR+CD38-cells, others suggest that HLA-DR+CD38- CD4+ T cells have a different functional profile than do HLA-DR+CD38+ cells in gut-associated lymphoid tissue in HIV (31). Thus, this relationship surfaced by VOLARE may inspire follow-up experiments.

**Figure 6.**
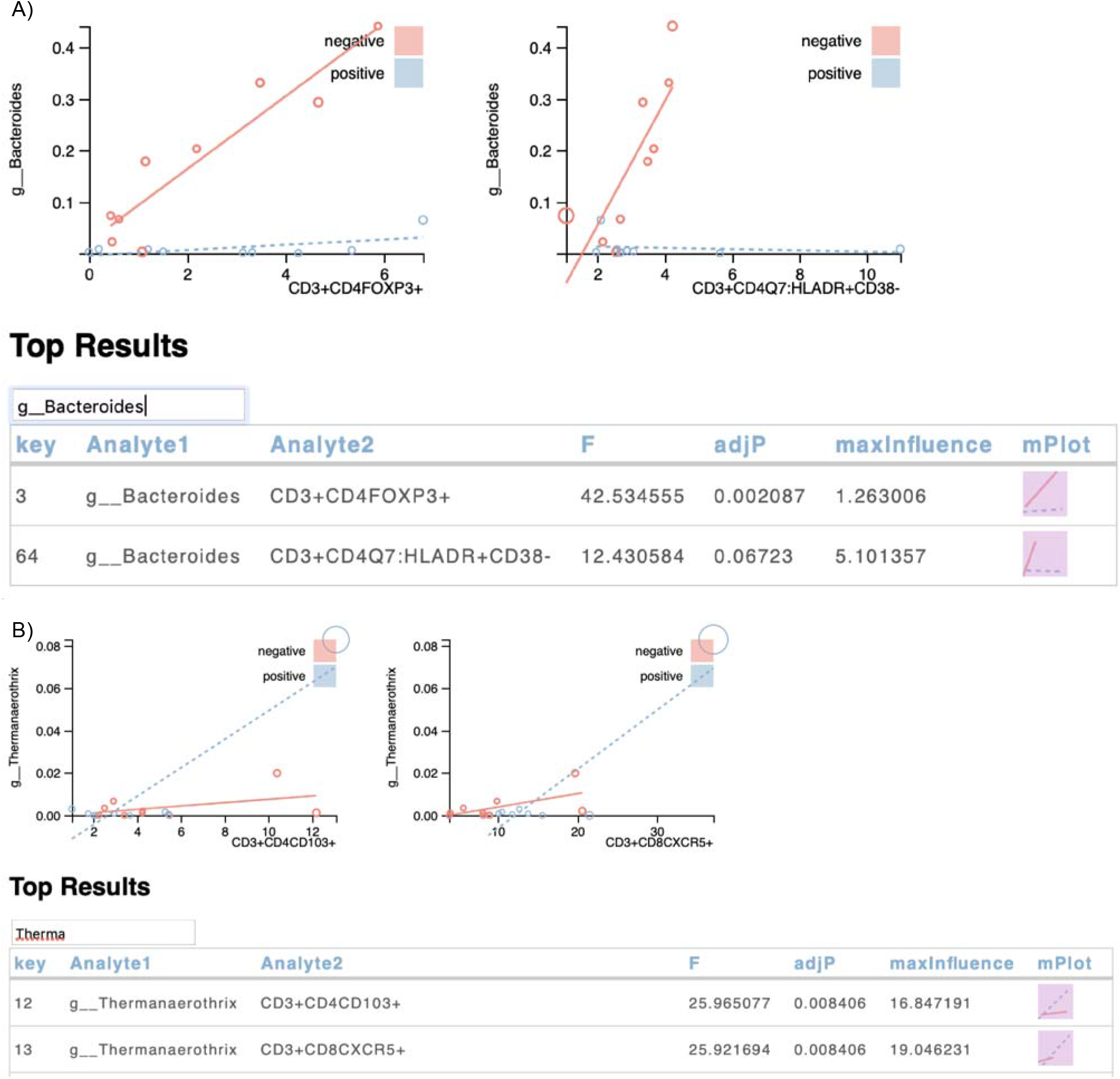
A) Two immune cell populations are strongly associated with *Bacteroides* in samples from HIV negative participants but not in HIV positive participants. The x-axis represents the percentage of the parent population (CD3+CD4+ T cells) that are FOXP3+ or HLADR+CD38-while the y-axis represents the microbiome data (measured in relative abundance, in the range from 0 to 1). Each point represents the values for one sample from one person. Points are color coded to represent the cohort to which the corresponding person belongs. Size of the points are proportional to the influence of the point on the fitted regression, with the maximum influence value shown in the top table. Lines represent the fitted regression model for each cohort. The closer the points are to the line, the better the model. B) Examples of relationships driven by an overly influential point found in the upper right hand corner of the detailed plots.

In this case study, the top table includes the largest absolute value of the difference in fits (DFFITS) metric (32) (labeled maxInfluence in the top tables in Figure 6), which enables domain experts to identify those relationships driven by an overly influential data point. DFFITS represents the number of standard deviations by which the i^th^ predicted value changes when the regression model is generated without the data for the i^th^ observation. Figure 6B illustrates two results in which there is one highly influential observation in the upper right-hand corner of the detailed plot. The radius of each circle is a function of the maximum influence, allowing visualization of highly influential observations in the context of the fitted model.

### Usage log analysis

Our users included six domain experts (two faculty members and one research assistant from the University of Colorado School of Medicine Division of Allergy and Clinical Immunology/Infectious Disease, one faculty member from the Division of Biomedical Informatics and Personalized Medicine, and one faculty member and one fellow from the Division of Rheumatology and Division of Gastroenterology, respectively) and two computational bioscience investigators. To better quantify usage patterns, we instrumented VOLARE to log user actions, such as loading files, searching the top table, and generating detailed plots. An analysis session might involve loading the same file several times to reset the visual display. Thus, we used the notion of an “analysis pass” to represent all of the activities from loading the file to the last action performed prior to resetting the display. Figure 7 illustrates metrics for 160 analysis passes collected over 55 days coming from 12 distinct IP addresses. The results show that most passes last ten minutes or less. The most common action in a pass is the generation of detailed plots, with an average of 12 plots per pass. Comparing the number of detailed plots generated per pass to the number of searches, we identified three main usage scenarios. One scenario is “big picture” generation of dozens of detailed plots, unaccompanied by searches. Another scenario is a mix of 2 to 5 searches and generation of 3 to 15 detailed plots (“search-inspect-search”). This may represent a cycle where one set of detailed plots leads the analyst to search for and inspect another set of detailed plots. A third scenario is zero or one searches combined with the generation of 1 to 5 detailed plots (“quick check”). This may represent a refinement of an earlier analysis, with a goal of generating a specific set of detailed plots for a screen capture, or a quick check of data. Taken together, these metrics illustrate that VOLARE supports a variety of exploration scenarios and that users are very interested in details-on-demand.

**Figure 7.**
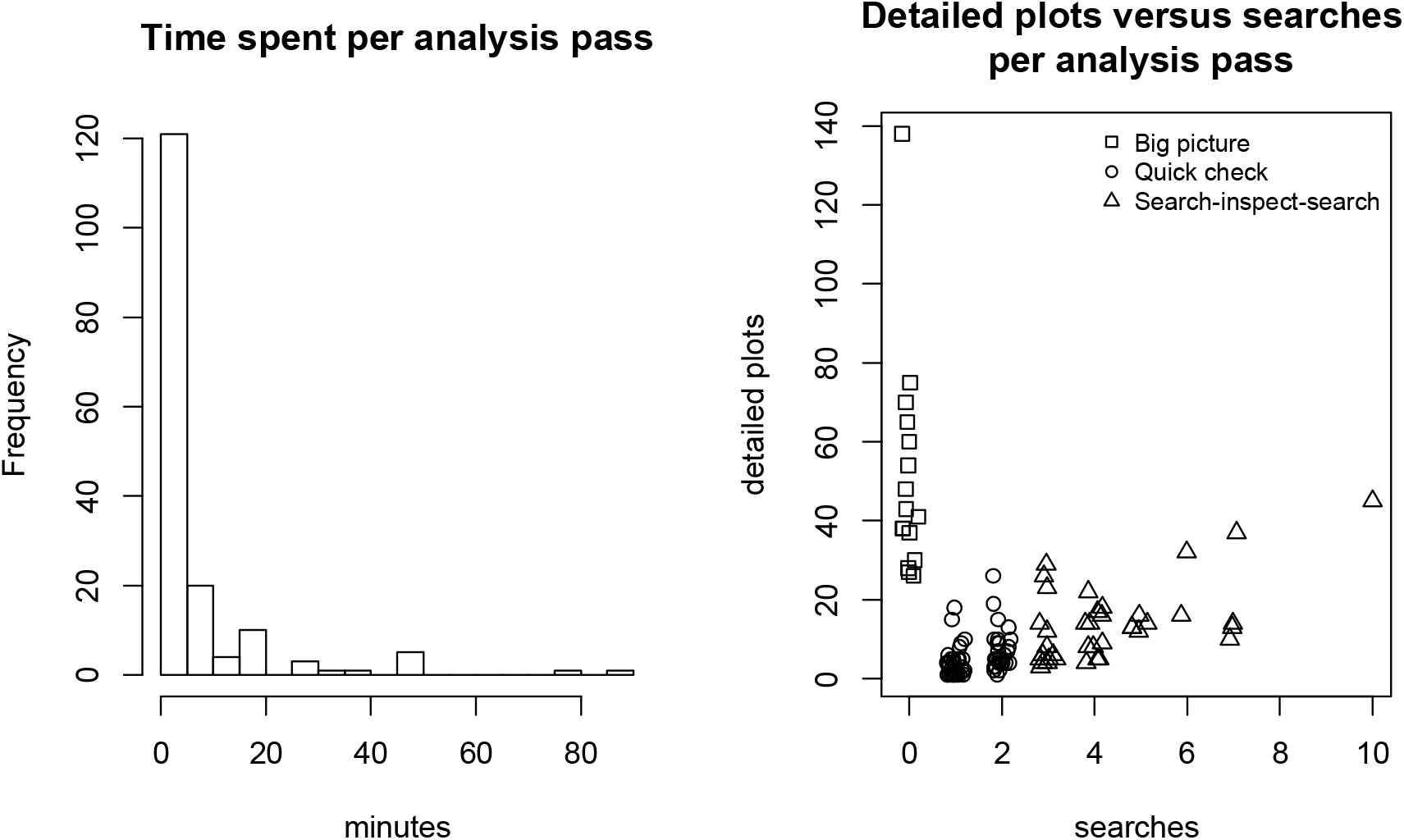
Usage scenarios. An analysis pass consists of loading a file and exploring the data, and lasts until the display is reset. (A) Most analysis passes last less than 20 minutes, but have lasted up to 90 minutes. (B) A comparison of the number of detailed plots generated versus the number of searches suggests three different analysis scenarios. One scenario is “big picture” generation of dozens of detailed plots, unaccompanied by searches (searches=0, dPlots greater than 20). Another scenario is a mix of 2 to 5 searches and generation of around 3 to 20 detailed plots (search-inspect-search). A third scenario is zero or one single searches combined with the generation of 1 to 10 detailed plots (quick check). Data is jittered on the horizontal axis to reduce overplotting.

## 4 Discussion

Across all three case studies, our domain experts had two main questions: (1) which microbes are differentially associated with which immune readouts by disease status; and (2) which of these candidates should we prioritize for follow-up laboratory experiments. To identify candidate relationships, we performed regressions across all microbe-immune readout pairs, while accounting for differences across cohorts. The regression framework supports an arbitrary number of cohorts and covariates such as age, sex, and study center; and offers established procedures for assessing statistical significance of various parts of the model, such as differences between cohorts. Given a top table of results, VOLARE aids domain experts in vetting these results. This vetting includes qualitative and quantitative assessments. Qualitative assessment considers the biological role of at least one of the analytes in the pair, and the ability to interrogate the relationship in an *in vitro* experiment. For example, if the microbe is readily available, the researcher can co-culture it with immune cells and measure immunological responses such as cytokine production, cell proliferation, and cell differentiation. Quantitative assessment considers both the magnitude of the readouts and the dynamic range of the relationships. The magnitudes should be large enough to be measured with precision, while the dynamic range should be large enough to be biologically meaningful.

To place VOLARE in context with existing visualization approaches, we consider three bodies of material: single assays, regression models, and biological networks. First, VOLARE complements existing approaches that support the visualization of the results of single assays such as 16S microbiome sequencing or CyTOF immunophenotyping. Our work is focused on identifying patterns across omes. As such, it differs from platform-specific tools, such as Qiime for analyzing 16S sequencing (36,37), and tSNE, SPADE, and Citrus for visualizing patterns in CyTOF data (38–40). These tools may perform feature extraction steps of identifying and quantifying analytes, be they sequences that have been assigned to a microbial taxonomy or clusters based on immune markers. Our work assumes such identification and quantification has been performed by a platform-appropriate pipeline. This allows us to focus on rich visual analysis tools that we can apply to a variety of omes. Second, Breheny and Burchett summarize over 40 years of work in visualization regression models in the introduction of their generalized approach for regression visualization, the R package visreg (41). Like them, we are focused on plotting models to illustrate fit. In general, visualizing model fit focuses on illustrating the results of a single regression model at a time. As such, there is limited emphasis on interactive visualization. In contrast, we consider dozens of fitted regression models concurrently. Siddiqui et al. integrated metabolomic and gene expression data using a linear model with an interaction term for phenotype (e.g. tumor versus non-tumor tissue in NCI-60 data sets (21)). The work described here differs in that we emphasize vetting of the results by domain experts. Third, approaches to biological network visualization are reviewed in (42). These interactive approaches tend to emphasize genomic relationships (e.g. genes and gene products, genes and transcription factors), supported by multiple lines of evidence, such as co-occurrence in a publication or pathway, or a straightforward experimental construct such as cell line:drug interaction (43). One of the challenges they tackle is filtering a very large number of relationships to a smaller, more manageable set that can be explored by a domain expert, as with RenoDOI (44). In contrast, rather than consider hundreds or thousands of relationships, we are focused on dozens. Navigating the “hairball” is less of a concern in this top table domain. Furthermore, in comparing the microbiome to immune cell repertoire, relationships may be speculative, and not yet catalogued in a reference database. The detailed regression plots allow the domain experts to assess the plausibility of these relationships.

There are several limitations to this work. First, as presented here, we have only considered two omes. While more omes could be included by increasing pairwise comparisons, the pairwise approach is self-limiting to a handful of omes. With two omes, there is one set of cross-omic pairwise comparisons; with three omes, three sets; with four omes, six sets; and in general, n(n-1)/2 sets, where n is the number of omes. That said, the support for visual analysis of promising results from the current two omes is a valuable contribution, setting the stage for extension more omes. Second, the regressions are performed by stand-alone computing resources, with necessary results and underlying details marshaled for the visualization layer. This means that changes to the regression model cannot be made on the fly in VOLARE. However, the regression analysis requires some statistical experience that the domain experts may not have. Thus, this is a natural breakpoint for separating the workflow. In addition, the existing approach of handing off data to a statistician for analysis has the same limitation. Third, we do not tune the regression model for each analyte pair. Instead, we use the same form of the regression model for all pairs, and support domain experts in vetting both model fit and biological relevance. Fourth, VOLARE does not provide strong support for extracting and modifying the graphs for presentation and publication. However, its main goal is data exploration rather than presentation. Currently, graphs can be extracted either by screenshot, or by copying svg elements from the document object model of the web page into an svg editor such as Inkscape or Adobe Illustrator.

Our future work includes adding features such as grouping by mPlot, searching the top table by Boolean expressions of analyte names, and displaying the detailed plot in response to clicking on a network edge. Grouping by mPlot would collect results that have similar association patterns across cohorts, such as a positive association in disease and a negative association in health. Searching by Boolean expressions of analyte names would enable domain experts to perform more powerful searches, such as “either of two specific microbial species combined with a particular immune cell activation marker.” Displaying the detailed plot in response to clicking on a network edge would lay the foundation for exploring paths of connected relationships. VOLARE can be applied to data sets that include different omics platforms, such as paired RNA-Seq and immune cell repertoire, and paired microbiome and metabolome; and to data sets that span more than two omes, such as microbiome, immune cell repertoire, and cytokine repertoire. We also plan to extend VOLARE to support regression models that may be more appropriate for microbiome data, such as the negative binomial (45).

## 5 Conclusion

VOLARE provides an interactive environment that that transcends the limitations of a static top table. Our usage analysis demonstrates that VOLARE supports a variety of analysis scenarios and that the detailed plots are an important component of user-driven analysis. We applied VOLARE to three case studies. In the fecal microbiome:cytokine study, we saw evidence of high IL-1α associated with high levels of a particular microbe, possibly suggesting an immune response. In the microbiome:IEL-produced cytokine study, we saw evidence of patient-level aberrations between several microbes and IL-6. In the gut biopsy microbiome:immune cell repertoire case study, we saw strong relationships between *Bacteroides* and both FOXP3+ CD4+ T cells and HLA-DR+CD38-CD4+ T cells in health but not in disease. VOLARE allows the domain expert to identify both patient-specific phenomena and relationships that are different by disease state. These relationships connect specific microbial taxa with specific immune system readouts, ideally at a level appropriate for follow-up experiments.

