## Supplemental Table 1 for "VOLARE: Visual analysis of disease-associated microbiome-immune system interplay"

Supplemental Table 1. Top table of microbe (Analyte1): cytokine (Analyte2) relationships from case study 1. F is from the partial F test of full model to reduced model, as described in Methods. P is the corresponding unadjusted p-value.

| Rank | Analyte1 | Analyte2 | F | p |
|---|---|---|---|---|
| 1 | Mb_3 | IL.22 | 52.2 | 1.19E-10 |
| 2 | Mb_15 | VEGF | 26.6 | 1.88E-07 |
| 3 | Mb_2 | ICAM.1 | 21.7 | 1.27E-06 |
| 4 | Mb_20 | VEGF | 21.2 | 1.55E-06 |
| 5 | Mb_8 | IL.15 | 19.0 | 4.04E-06 |
| 6 | Mb_8 | IL.1alpha | 16.5 | 1.29E-05 |
| 7 | Mb_15 | IL.12.23p40 | 15.7 | 1.98E-05 |
| 8 | Mb_2 | IL.12.23p40 | 14.9 | 2.97E-05 |
| 9 | Mb_8 | IL.16 | 14.0 | 4.76E-05 |
| 10 | Mb_7 | IL.7 | 13.4 | 6.48E-05 |
| 11 | Mb_6 | IL.1alpha | 11.7 | 1.65E-04 |
| 12 | Mb_4 | Calprotectin | 10.7 | 2.97E-04 |
| 13 | Mb_8 | VEGF | 10.6 | 3.11E-04 |
| 14 | Mb_22 | IL.22 | 10.2 | 3.85E-04 |
| 15 | Mb_8 | TNF.b | 9.8 | 4.93E-04 |
| 16 | Mb_8 | GM.CSF | 9.0 | 8.18E-04 |
| 17 | Mb_22 | sIgA | 8.2 | 1.41E-03 |
| 18 | Mb_21 | IL.15 | 8.2 | 1.41E-03 |
| 19 | Mb_15 | IL.16 | 7.9 | 1.64E-03 |
| 20 | Mb_20 | IL.15 | 7.5 | 2.18E-03 |
| 21 | Mb_10 | IL.16 | 7.1 | 2.81E-03 |
| 22 | Mb_2 | TNF.b | 7.1 | 2.97E-03 |
| 23 | Mb_2 | VCAM.1 | 7.0 | 3.10E-03 |
| 24 | Mb_20 | Calprotectin | 6.9 | 3.25E-03 |
| 25 | Mb_5 | IL.22 | 6.9 | 3.36E-03 |
| 26 | Mb_2 | GM.CSF | 6.8 | 3.64E-03 |
| 27 | Mb_17 | IL.1alpha | 6.7 | 3.79E-03 |
| 28 | Mb_14 | IL.12.23p40 | 6.7 | 3.80E-03 |
| 29 | Mb_19 | sIgA | 6.4 | 4.63E-03 |
| 30 | Mb_11 | VCAM.1 | 6.4 | 4.70E-03 |
| 31 | Mb_14 | VEGF | 6.4 | 4.70E-03 |
| 32 | Mb_15 | IL.15 | 5.6 | 8.68E-03 |
| 33 | Mb_2 | IL.7 | 5.4 | 9.53E-03 |
| 34 | Mb_21 | VEGF | 5.2 | 1.15E-02 |
| 35 | Mb_14 | ICAM.1 | 5.1 | 1.21E-02 |
| 36 | Mb_16 | sIgA | 5.0 | 1.34E-02 |
| 37 | Mb_8 | IL.12.23p40 | 4.6 | 1.84E-02 |
| 38 | Mb_15 | ICAM.1 | 4.5 | 1.99E-02 |

| | | | | |
|---|---|---|---|---|
| 39 | Mb_4 | CRP | 4.3 | 2.18E-02 |
| 40 | Mb_20 | IL.12.23p40 | 4.3 | 2.29E-02 |
| 41 | Mb_1 | CRP | 4.2 | 2.37E-02 |
| 42 | Mb_2 | IL.22 | 4.1 | 2.60E-02 |
| 43 | Mb_12 | CRP | 4.0 | 2.81E-02 |
| 44 | Mb_9 | sCD14 | 3.8 | 3.38E-02 |
| 45 | Mb_13 | Calprotectin | 3.7 | 3.60E-02 |
| 46 | Mb_17 | sIgA | 3.7 | 3.66E-02 |
| 47 | Mb_10 | IL.1alpha | 3.7 | 3.69E-02 |
| 48 | Mb_4 | VEGF | 3.5 | 4.15E-02 |
| 49 | Mb_13 | IL.1alpha | 3.5 | 4.24E-02 |
| 50 | Mb_4 | ICAM.1 | 3.5 | 4.28E-02 |
| 51 | Mb_12 | Calprotectin | 3.5 | 4.30E-02 |
| 52 | Mb_20 | IL.16 | 3.5 | 4.38E-02 |
| 53 | Mb_12 | IL.1alpha | 3.4 | 4.47E-02 |
| 54 | Mb_8 | Calprotectin | 3.4 | 4.51E-02 |
| 55 | Mb_4 | IL.1beta | 3.4 | 4.53E-02 |
| 56 | Mb_7 | IL.1alpha | 3.4 | 4.80E-02 |
| 57 | Mb_18 | IL.16 | 3.4 | 4.81E-02 |
| 58 | Mb_3 | sIgA | 3.3 | 4.86E-02 |
