## Supplemental File 1 for "VOLARE: Visual analysis of disease-associated microbiome-immune system interplay"

**Supplemental File 1. Biological methods and materials**

***Study participants, case studies 1 and 3***

Study participants in case studies 1 and 3 were recruited as described in Armstrong et al (1). All subjects were recruited from the Denver, Colorado metropolitan area. For the high-risk men having sex with men in Case study 1, high risk for HIV infection was defined as in a prior study of a candidate HIV vaccine: (1) a history of unprotected anal intercourse with one or more male or male-to-female transgender partners, (2) anal intercourse with two or more male or male-to-female transgender partners, or (3) being in a sexual relationship with a person who has been diagnosed with HIV (2). The HIV positive individuals in case study 3 had chronic HIV-1 infection and were on Antiretroviral Therapy (ART) for ≥ 12 months with a minimum of three ART drugs prior to study entry and < 50 copies HIV RNA/mL within 30 days prior to study entry and no plasma HIV-1 RNA ≥ 50 copies/mL in the past 6 months. Individuals who reported taking antibiotics within 3 months of sample collection were excluded from the study. The fecal microbiome data from these subjects were previously described (1).

***Microbiome sample collection and quantification, case study 1***

Fecal swabs were collected from study participants. DNA extracted using the PowerSoil Kit from the gut biopsies were subjected to 16S rRNA sequencing. The V4 region of 16S rRNA was targeted using protocols of the Earth Microbiome Project (3). Sequences were denoised with DADA2 (4), and assigned to genera using the RDP classifier trained on the greengenes 13_8 dataset using QIIME 2(5).

***Fecal water preparation and quantification, case study 1***

Fecal water was generated from frozen feces by first combining it with a saline solution (DPBS, Protease Inhibitor, EDTA, DNase).  Samples were homogenized, centrifuged at high speed and the supernatant was filtered, and stored at -80˚C until testing.  Standard sandwich ELISAs and Multi-plex ELISAs (MesoScale Diagnostics, Rockville, MD) were used to measure cytokines, chemokines and markers of gut epithelial integrity.

***Microbiome sample collection and quantification, case study 3***

Colonic biopsies were collected by flexible sigmoidoscopy from study participants. Flash frozen biopsies were homogenized using bead beating. DNA extracted using the PowerSoil Kit from the gut biopsies were subjected to 16S rRNA sequencing. The V4 region of 16S rRNA was targeted using protocols of the Earth Microbiome Project Project (3). Sequences were denoised with DADA2 (4), and assigned to genera using the RDP classifier trained on the greengenes 13_8 dataset using QIIME 2(5).

***Immunophenotyping sample collection and quantification, case study 3***

Colonic biopsies, as described above, were homogenized to single cell suspension following enzymatic digestion. Single cells were filtered and subjected to surface and intracellular staining with 35 metal-tagged antibodies. Cells were analyzed by a CyTOF 2 and cell populations were determined using FlowJo software.

***Case study 2***

Study participants and sample collection are described in Regner et al. (6)
